# Disentangling the Relative Strength of Niche Competition from Grazing-Induced Phytoplankton Mortality

**DOI:** 10.1101/072231

**Authors:** Stephen J. Beckett, Joshua S. Weitz

## Abstract

The dilution method is the principal tool used to infer *in situ* microzooplankton grazing rates. However, grazing is the only mortality process considered in the theoretical model underlying the interpretation of dilution method experiments. Here we evaluate the robustness of mortality estimates inferred from dilution experiments when there is concurrent niche competition amongst phytoplankton. Using a combination of mathematical analysis and numerical simulations, we find that grazing rates may be overestimated – the degree of overestimation is related to the importance of niche competition relative to microzooplankton grazing. In response, we propose a conceptual method to disentangle the effects of niche competition and grazing by diluting out microzooplankton, but not phytoplankton. Our theoretical results suggest this revised “Z-dilution” method can robustly infer grazing mortality, regardless of the dominant phytoplankton mortality driver in our system. Further, we show it is possible to independently estimate both grazing mortality and niche competition if the classical and Z-dilution methods can be used in tandem. We discuss the significance of these results for quantifying phytoplankton mortality rates; and the feasibility of using the Z-dilution method in practice.

## Introduction

Phytoplankton form the base of the ocean food web and are drivers of ocean biogeochemical cycles. Microzooplankton grazing is thought to be one of the dominant drivers of phytoplankton mortality [1, 2]; and is a core process within the marine microbial loop [3, 4]. However, other processes such as nutrient limitation, sinking and viral lysis compete and interact with grazers as sources of phytoplankton mortality [5–8]. Understanding the dynamics of microbial food webs is therefore key to understanding their role in oceanic biogeochemical fluxes [9–11].

Estimates of the relative importance of grazing versus other mortality drivers depends on the quality and robustness of experimental techniques. The dilution method [12] is a long-established and popular technique used to measure the impact of micro- and nano-zooplankton on phytoplankton communities. The method is outlined in Fig 1. A sample of seawater is taken and prefiltered so only the microbial fraction remains. Some of this whole seawater (WSW) sample is further filtered, to remove phytoplankton and microzooplankton, creating a diluent. The dilution method procedure creates a series of bottles each containing a different proportion of WSW mixed with diluent to create a dilution series.

**Fig 1.**
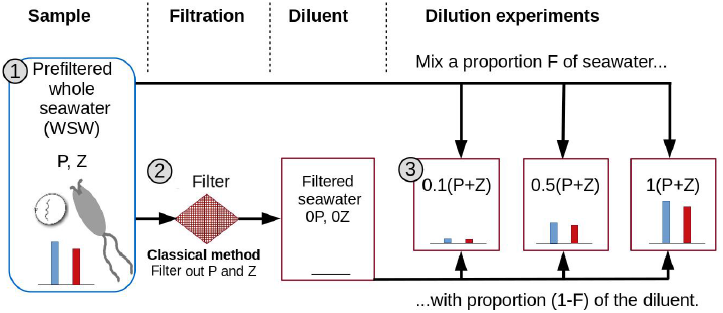
Schematic of the classical dilution method of Landry and Hassett. **1. Sample** Environmental samples are prefiltered to focus on microbial communities - this is termed whole seawater (WSW). Dilution method theory assumes WSW contains microzooplankton and the phytoplankton they graze upon. **2. Filtration** The classic dilution method filters some WSW to create a diluent containing no phytoplankton or microzooplankton. **3. Dilution series** A series of bottles are filled with a proportion *F* of WSW and mixed with a proportion (1 − *F*) of the diluent creating a dilution series. The blue and red bars represent the relative abundance of phytoplankton and microzooplankton. Apparent growth rates are calculated by measuring the differences in phytoplankton population sizes in each bottle across the dilution series at two time points (the beginning and end of an incubation period). The microzooplankton grazing rate is estimated by finding the gradient of a linear regression model between the dilution level *F* and the apparent growth rate.

By measuring the differences in phytoplankton population sizes between two time points within each bottle, the corresponding approximate per capita growth rates can be calculated. The dilution curve represents a plot of the approximate growth rates within each bottle against the dilution level (the proportion of WSW within a bottle). The phytoplankton growth rate and the grazing rate are reported as the intercept and slope, respectively, of the dilution curve. This is due to the underlying theoretical model of phytoplankton dynamics presumed to operate in the bottle. This model, introduced by Landry and Hassett [12], is:

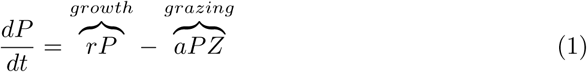
in which *P* is the density of phytoplankton, which grow at rate *r*; and are grazed upon by microzooplankton *Z* at a rate of *a*. Consider that the whole seawater sample initially contains a phytoplankton density of *P*_0_ and a microzooplankton density of *Z*_0_. A particular bottle within the dilution series containing a proportion *F* of whole seawater will initially contain phytoplankton and microzooplankton densities of *FP*_0_ and *FZ*_0_ respectively. Evaluating the apparent growth rate, calculated as the rate of population change per capita, under these conditions shows that:

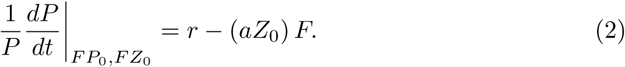

Eq (2) implies that the relationship between dilution level and apparent growth rate will be linear with an intercept equal to the intrinsic per capita growth rate and a gradient equal to the initial per capita grazing mortality rate. However, the model given by Eq (1) may be an insufficient description of bottle phytoplankton dynamics for several reasons.

Microzooplankton grazing may respond non-linearly with respect to phytoplankton densities in the bottle for example due to feeding saturation. This could significantly alter the shape of the dilution curve and how grazing mortality estimates can be made [13–15]. Additional sources of phytoplankton mortality such as losses due to sinking or viral lysis (e.g. [16,17]) are not accounted for in this model, yet incorporating these processes may be important for interpreting empirical data from dilution experiments. We also note that the model intends to describe bulk community dynamics. Diversity is not considered as phytoplankton and microzooplankton communities are each treated as a single population, neglecting potential important functional differences between species. Calbet et al. [18] consider trophic chains; but overall limited attention has been given to diversity.

Another critique of the method is that it does not take into account the resource requirements for phytoplankton growth. In the absence of microzooplankton (*Z*_0_ = 0) this model predicts the phytoplankton population will grow without bounds to infinity as niche competition between phytoplankton is not considered. Subsequent modifications of the dilution method included a nutrient enrichment step so that phytoplankton could grow near their idealized maximum rates [14]. Even if nutrients are added to bottles to attempt to keep phytoplankton growing in the exponential growth phase, this does not eliminate the potential of competition occurring between phytoplankton. While many potential limitations exist (some of which are included in the discussion), in this paper we choose to focus on how the inclusion of niche competition may alter the interpretation of dilution experiment measurements.

A common way to represent niche competition between phytoplankton is by using a logistic growth model [19]. Logistic growth is a phenomenological model used to implicitly represent competition for resources, whilst not tracking those resources explicitly. It is a common tool in mathematical ecology and has been used to describe phytoplankton growth dynamics (e.g. [20]). Using a logistic growth function has the effect of bounding phytoplankton populations to a carrying capacity *K*. We note that the inclusion of logistic growth could be interpreted to mean that phytoplankton mortality increases as *P* approaches *K* or that phytoplankton growth decreases as *P* approaches *K*, or a combination thereof. A candidate model considering the effect of niche competition is:

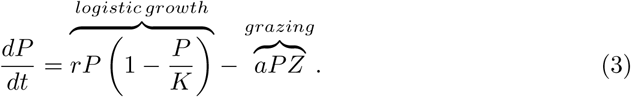

Using Eq (3) as the description for phytoplankton dynamics we find the apparent growth rate within a bottle in the dilution series is predicted as:

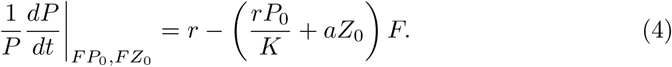

This model, including niche competition, also predicts a linear relationship between dilution level and apparent growth rates. Here, the intercept of the dilution curve represents the maximum growth rate, just as in the classical model. However, the slope is now interpreted as the *combined* effect of niche competition and grazing, both of which necessarily have positive signs. Therefore, dilution experiments may overestimate grazing rates should niche competition be important. We hypothesize that niche competition will be important when the functional form of phytoplankton growth is better approximated by a logistic growth function than by an exponential growth function. In this paper we explore this hypothesis and ways to improve dilution experiments when both grazing and niche competition operate concurrently.

## Materials and Methods

In this paper we utilize ecological models of phytoplankton dynamics analyzed in silico. For the *in silico* dilution experiments, whole seawater is mixed with the diluent at 10 dilution levels (with proportions *F* = 0.1, 0.2, 0.3, 0.4, 0.5, 0.6, 0.7, 0.8, 0.9 and 1 of WSW) to create a dilution series with 10 bottles. The apparent growth rate,
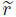, within each bottle is calculated from recordings of the simulated population dynamics as:

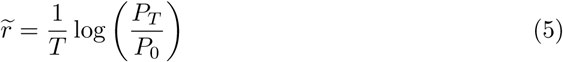
where *P*_0_ is the phytoplankton density at the beginning of the simulation and *P*_*T*_ is the phytoplankton density after time *T* [12]. Here the apparent growth rate is calculated using an incubation time of 24h in length. Dilution method estimates of grazing mortality are then calculated by finding the slope of the linear regression fit of apparent growth rates within each simulated bottle in the dilution series (response) against the proportion, *F*, of WSW it contains (predictor). The intercept of this regression is understood as representing phytoplankton growth rate, whilst the slope is interpreted as the grazing mortality rate. Model parameters for *in silico* dilution experiments are shown in Table 1. Code for the following analyses is available [21].

**Table 1.** Ecological parameters used in this study.

| Symbol | Description | Value | Units |
| --- | --- | --- | --- |
| $r$ | Intrinsic per capita growth rate of phytoplankton | 1 | day<sup>-1</sup> |
| $K$ | phytoplankton carrying capacity | $2.2 \times 10^7$ | phytoplankton ml<sup>-1</sup> |
| $a$ | microzooplankton attack rate | $4.8 \times 10^{-5}$ | ml/(microzooplankton·day) |
| $Z_{0L}$ | Initial microzooplankton density at low pressure | 1000 | microzooplankton ml<sup>-1</sup> |
| $Z_{0I}$ | Initial microzooplankton density at intermediate pressure | 10000 | microzooplankton ml<sup>-1</sup> |
| $Z_{0H}$ | Initial microzooplankton density at high pressure | 20000 | microzooplankton ml<sup>-1</sup> |
| $P_0$ | Initial phytoplankton density | $K \left(1 - \frac{aZ_0}{r}\right)$ | phytoplankton ml<sup>-1</sup> |

## Assessment

### Top-down pressure as indicator of control of phytoplankton by grazers

In the absence of microzooplankton the steady state solution for phytoplankton population density within the bottle is 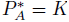. Similarly, the steady state in the presence of microzooplankton is 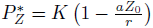. Neither of these densities are necessarily reached during the dilution experiment. Nonetheless, these densities provide a means to quantify the relative importance of grazing. To see why, note that increasing microzooplankton pressure will lead to a reduction in 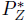, away from 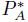.

Microzooplankton pressure is maximised when the per capita mortality rate due to grazing, *aZ*_0_, is equal to the per capita intrinsic phytoplankton growth rate, *r*. We can quantify top-down control of phytoplankton by the microzooplankton by using *δ_Z_* as a measure of **grazing pressure**:

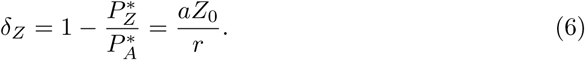

In the absence of microzooplankton, no grazing occurs and *δ_Z_* = 0. If top-down microzooplankton grazing were to drive steady state phytoplankton density to half of the resource limited density then *δ_Z_* = 0.5. When microzooplankton grazing results in the phytoplankton population being drawn to extinction 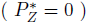, then top-down control by microzooplankton is maximised and maximised and *δ_Z_* = 1. We use three specific levels of grazing pressure (**low:** *δ_Z_* = 0.048, *aZ*_0_ = 0.002, **intermediate:** *δ_Z_* = 0.48, *aZ*_0_ = 0.02, **high:** *δ_Z_* = 0.96, *aZ*_0_ = 0.04) to highlight the performance and sensitivity of the dilution method to the dominance of microzooplankton grazing as a driver of phytoplankton mortality.

### Sensitivity of mortality rate estimates to niche competition

The ability of the dilution method to estimate microzooplankton associated mortality rates is potentially affected by the relative level of bottom-up (i.e., niche competition) and top-down (i.e., microzooplankton) pressures. This is highlighted in Fig 2A using simulations with the three chosen levels of grazing pressure. Each level of grazing pressure corresponds to a steady state microzooplankton density (see Table 1) from which a steady state phytoplankton density, *P*\*, is calculated while keeping the growth rate, *r*, fixed at 1 per day. The grazing mortality rate was estimated using the dilution experiment, in order to compare to the baseline mortality rate *aZ*_0_. Estimates made using the classical dilution method were closest to the baseline rates when grazing pressure was high, but substantially higher than the expected baseline rates when grazing pressure was low. Mortality rate bias is calculated as the estimated grazing mortality rate (by the dilution method) divided by the baseline grazing mortality rate. Examining the resulting bias in mortality rates (Fig 2B) we find that baseline mortality is overestimated by a factor of ≈ 20 at the highlighted low grazing pressure, a factor of ≈ 2 at the highlighted intermediate grazing pressure and by less than 5% at the highlighted high grazing pressure. Thus mortality rate bias is lowest when grazing pressure is highest, corresponding to situations where phytoplankton densities approach 0. This suggests that the classical dilution method provides more accurate estimates of microzooplankton associated mortality rates when grazing pressure is high i.e., *when grazing is significantly more important than niche competition.*

### Conflating grazing with population growth inhibition

Grazing mortality rates are overestimated by the classical dilution method when microzooplankton grazing pressure is low. Indeed, the estimated grazing mortality rates appear closer to the maximum phytoplankton growth rate (see Fig 2A). This feature of the classical dilution method arises due to the inability of the method to disentangle grazing and niche competition as shown in Eq (4). As a simple modification, we assume that the WSW sample is dynamically at steady state; *P*\* and *Z*\* denoting the steady state densities in the environment. Therefore, the initial conditions for our *in silico* experiments for a bottle with dilution factor *F* are *P* = *FP*\* and *Z* = *FZ*\*, where *Z*\* = *Z*_0_ and 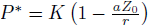. Substituting these quantities into Eq (4) we find:

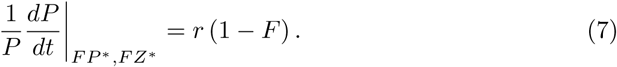

**Fig 2.**
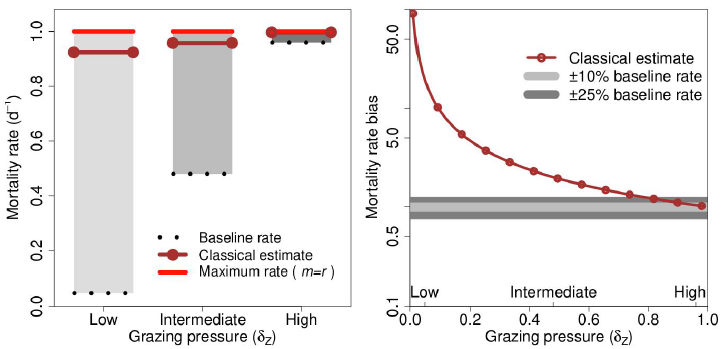
The classical dilution method may overestimate rates of mortality via grazing. (A) Expected baseline microzooplankton associated mortality rates and rates estimated using the classical dilution method for three levels of grazing pressure; low grazing pressure (1000 microzooplankton ml^−1^), intermediate grazing pressure (10000 microzooplankton ml^−1^) and high grazing pressure (20000 microzooplankton ml^−1^). The maximum mortality rate is calculated for the condition when total mortality, *m*, is equal to the phytoplankton growth rate *r*. (B) Mortality rate bias across the full gradient of grazing pressure. The grazing pressure associated with each of the examples given in (A) are shown on the x-axis.

In the limit when *F* = 0, we find that the per capita rate of change is *r*; whilst at *F* = 1, the per capita rate of change is 0. At steady state phytoplankton growth is balanced by phytoplankton mortality. When microzooplankton grazing pressure is high, mortality due to grazing is much greater than mortality due to niche competition and so *r* ≈ *aZ*_0_. However, as grazing pressure decreases the relative importance of mortality due to competition between phytoplankton increases and *r* is no longer a good estimate of grazing mortality. Thus, an alternative interpretation for the slope calculated by the classical dilution method is that it represents **the combined rate of phytoplankton mortality by both microzooplankton grazing and by niche competition.**

### Diluting microzooplankton alone enhances grazing rate estimates

In the classical dilution method the incubated samples contain diluted levels of both phytoplankton and microzooplankton [12]. Rather than enriching the medium (see [14]) we propose a different approach: altering the filter to exclude microzooplankton but not phytoplankton cells. Thus, altering the proportion of WSW used within each treatment will only change the initial microzooplankton density – representing a linear gradient between no microzooplankton and ambient levels of microzooplankton; whilst maintaining ambient levels of phytoplankton. We first explore this approach conceptually and revisit the practical constraints of implementation in the Discussion. This “Z-dilution” method, is depicted in Fig 3.

**Fig 3.**
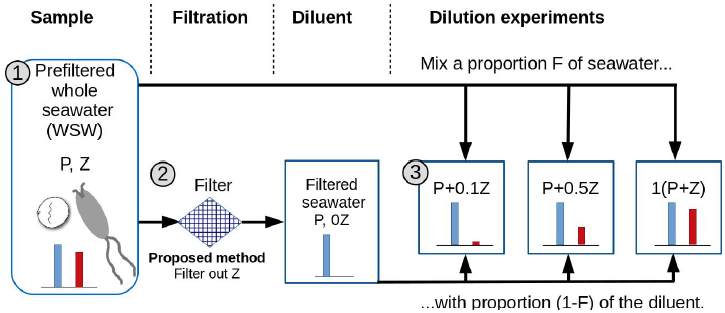
Proposed revision to the classical dilution method. Whilst the classical dilution method (see Fig 1) uses a filter excluding phytoplankton and microzooplankton, the proposed method instead uses an alternative filter, able to exclude microzooplankton, but through which phytoplankton can pass. Thus constituent levels of microzooplankton and phytoplankton within each bottle, shown by red and blue bars respectively, differ to those in the classical dilution experiment.

In this case, we expect the per capita bottle population dynamics for a dilution level *F* to be:

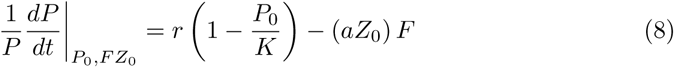
where the slope of the dilution curve is here found as the grazing rate. This slope is precisely the baseline grazing mortality rate we hope to estimate using the dilution method. This analytical result suggests this revised method should accurately estimate grazing mortality rates, regardless of the level of grazing pressure.

As before, we can investigate the expected per capita growth rate *in silico* using the Z-dilution method under steady state dynamics. By substituting the initial conditions for this revised dilution method, *P* = *P*\* and *Z* = *FZ*\*, into Eq (4) we obtain the per capita phytoplankton rate of change as:

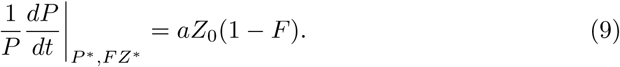

Under steady state conditions we expect the dilution curve to have a slope of *aZ*_0_ and an intercept of *aZ*_0_.

### Comparison of classical and revised dilution methods

We now compare the performance and robustness of the classical and Z-dilution methods. We do so by varying the strength of grazing pressure. Fig 4 shows the dilution curves measured for both methods after 24h incubation for the three highlighted conditions of grazing pressure. The linear regression fits for the classical dilution method have a slope (and intercept) close to the intrinsic growth rate under all three conditions. In contrast, the slope (and intercept) found by the Z-dilution method is closer to the identified baseline rates of microzooplankton associated mortality rate. This provides confirmatory evidence to support the analytical results found in Eq (7) and Eq (9).

**Fig 4.**
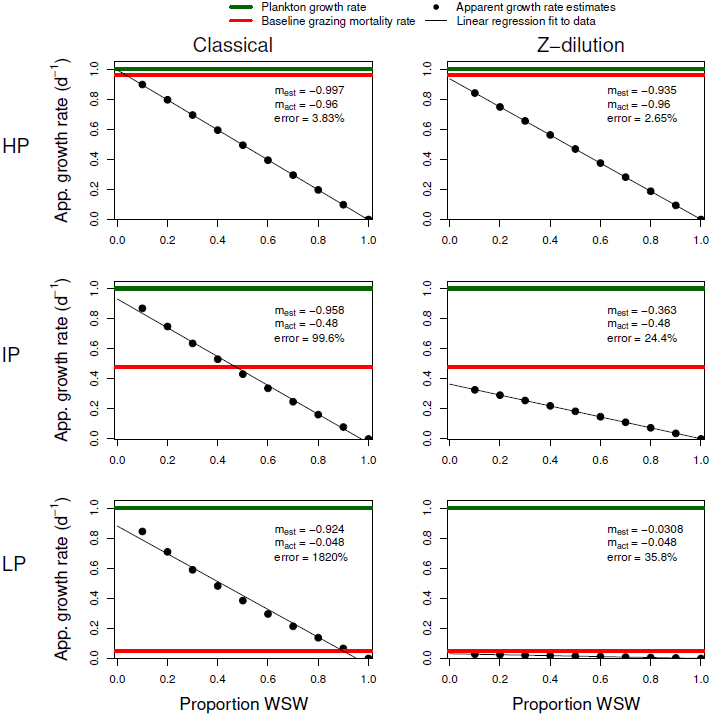
Dilution curves show the classical dilution method is insensitive to niche competition. Apparent growth rates are plotted against the proportion of whole seawater for each bottle in the *in silico* dilution series after a 24h incubation period when using the classical dilution method and the Z-dilution method respectively. Three cases, each with different microzooplankton grazing pressure conditions (LP: Low pressure, IP: Intermediate pressure and HP: high pressure, as defined in Fig 2) are shown. The estimated mortality rate (m_est_) found as the linear regression slope, the baseline mortality rate (m_act_) and the percentage error in estimation are shown for each subplot (all rounded to 3 s.f.).

As a consequence, the classical and Z-dilution methods give substantially different estimates of grazing mortality rates (Fig 5). When microzooplankton grazing pressure is high estimates made by both methods are within 10% of the true mortality rate after a 24h incubation period. However, the ability to estimate mortality rates declines with reduced grazing pressure. The classical dilution method overestimates microzooplankton associated mortality. The revised method, in which only microzooplankton are subject to dilution, tends to underestimate the true rate. The degree to which the revised method underestimates the expected rates is substantially less than that by which the classical dilution method overestimates the expected rates. This is particularly evident when grazing pressure is low.

**Fig 5.**
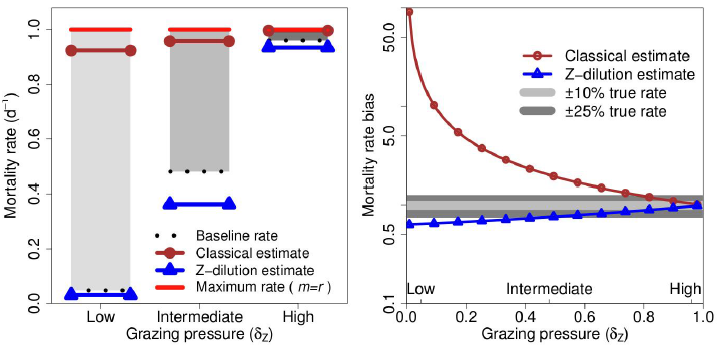
A comparison of classical and Z-dilution method estimates. (A) Mortality rates and their estimates at three levels of grazing pressure after 24h incubation period. The maximum mortality rate is calculated for the condition when the mortality, *m*, is equal to the phytoplankton growth rate *r*. Baseline mortality rates are shown for each condition. (B) Mortality rate bias is plotted against the level of grazing pressure (*δ_Z_*) - the three conditions shown in (A) are marked on the x-axis. Bands indicating ±10% and ±25% differences from the true mortality rate in the sample are shown.

## Discussion

Estimated microzooplankton grazing rates are central to efforts to understand the relative importance of top-down vs. bottom-up forces in the global oceans. We find that the performance of mortality rate estimates is dependent on the filtering apparatus used by the dilution method and to the relative amount of top-down control. In this paper we reviewed the robustness of the classical dilution method and a proposed alternative, the “Z-dilution” method. Classical dilution theory works well when niche competition is low relative to microzooplankton grazing. But, in circumstances when niche competition is an important control of phytoplankton populations, we predict that the classical method will lead to an overestimation of grazing mortality. Instead, we find that diluting microzooplankton, but not phytoplankton, represents a conceptually feasible approach to isolate the effects of grazing from niche competition. In doing so, the method goes beyond efforts to estimate grazing exclusively - whether in the classical dilution method or in extensions that include enrichment [12, 14].

We found that the slope of the dilution curve in the classical dilution method is expected to be equivalent to the combined effect of niche competition and microzooplankton grazing (Eq (4)). Our work suggests that if the actual population dynamics within each dilution bottle includes both grazing mortality and niche competition, then finding the differences between the slopes (or equivalently, using this model, the intercepts) of the classical and the Z-dilution methods would provide a way to quantify the effects of niche competition. We note that an alternative and complementary approach for assessing the relative impact of niche competition could be found by measuring the differences between the slopes of the classical [12] and nutrient-enriched [14] dilution experiments. If niche competition is unimportant, then one should expect to recover the same dilution curve for the classical, the nutrient-enriched, and the Z-dilution methods. We also note that in the Z-dilution method we assume the use of a filter that can exclude microzooplankton, but allow all phytoplankton to pass through. In practice, it will be important to evaluate the extent to which size- or feature-specific filtering can be achieved as they may share similar and overlapping size-spectra. This may limit application of the Z-dilution method to well-characterized artificial or lab ecosystems where an appropriate filtering procedure can be chosen, unless novel filtering methods can be developed or further theoretical work can demonstrate the robustness of the Z-dilution method to partial overlap. We encourage empiricists to develop model systems with which the Z-dilution methodology and its corresponding analysis may be validated, such that the relative strength of niche competition can be evaluated. This could be achieved by choosing a grazing species whose size-spectra does not overlap with that of its prey.

This is not the first time that the use of the dilution method has been critically assessed (e.g. [13–15, 18, 22]) or suggested to overestimate the impact of grazing. Indeed, a recent meta-analysis found that many studies erroneously applied tests that assessed the significance of mortality rates based on whether they were greater than or less than zero, rather than strictly greater than zero [23, 24]. Other studies have also suggested that grazing rates may be overestimated due to changes within the grazer communities [25]. In addition to statistical and measurement uncertainty it is important to address model uncertainty. As populations are only measured twice during dilution experiments, pre- and post-incubation, it is unknown to what extent the assumption of exponential growth given by classical dilution theory is held. Assuming grazer communities are constant on the timescale of dilution experiments we found that microzooplankton associated mortality rates, found via the classic dilution method, may be overestimated in environments when microzooplankton grazing pressure is low. However, we suggest that previous rate estimates made by the classical dilution method could potentially be used as an upper bound for the true grazing mortality rate. Similarly, estimates made via nutrient-enhanced approaches could be compared to those here as a means to gauge the *relative* importance of grazing with respect to niche competition.

The model presented in our manuscript is purposefully simple in order to convey our key message: classical grazing rate estimates may be conflated with niche competition. The ability to which it is possible to critically measure ecological properties using dilution experiments hinges on the appropriate formulation of the underlying ecological model used to interpret the data. Future theory work should focus on more realistic model descriptions of phytoplankton interactions and dynamics. Investigating community diversity e.g. differences in prey selectivity, allelopathic competition, virus host range and other important ecological traits, could be important for accurately interpreting the bulk community rates obtained by performing dilution experiments [26]. Another important focus could be the relevance of mixotrophic lifestyles [27, 28] to dilution rate measurements. We encourage those who use the dilution method to investigate the Z-dilution method as a way to gain quantitative understanding of the importance of niche competition. Additionally, we encourage researchers to investigate the response of other components of phytoplankton mortality with respect to dilution. The present analysis further supports the need to combine theory and experiments together to improve understanding of ecosystem functioning of marine microbes.

## Acknowledgments

This work was supported by a grant from the Simons Foundation (SCOPE Award ID 329108, J. S. W.). We thank David Caron, Debbie Lindell, Mick Follows and anonymous reviewers for feedback that has improved this manuscript.

